# A GreenGate-compatible vector set for efficient protein purification from E. coli

**DOI:** 10.64898/2026.06.12.731827

**Authors:** Maria Dolores López-Bueno, Franziska Anzenberger, Andrea Lepper, Andrea Bleckmann, Philipp Denninger

**Author notes:** Corresponding author: Philipp Denninger. These authors contributed equally (Maria Dolores López-Bueno, Franziska Anzenberger, Andrea Bleckmann, Philipp Denninger).

## Abstract

Recombinant protein purification from *E. coli* frequently requires screening various affinity tags and variations to optimize yield and purity. However, classical cloning methods are limited in throughput and modularity, which is circumvented by GoldenGate cloning. While GreenGate cloning, a GoldenGate variant, is widely used in plant research, it lacks compatibility with *E. coli* expression vectors. Here, we introduce a comprehensive, GreenGate-compatible vector toolkit for efficient and versatile assembly of three modules, for N- and C-terminal tagging of a protein of interest into an IPTG-inducible *E. coli* expression vector. To allow versatility, this toolkit contains diverse affinity tags, with or without HRV3C protease cleavage sites. Moreover, we included plasmids for the homemade low-cost production of this protease. This GreenGate-compatible toolkit allows efficient combinations of different tags and eliminates the need for re-cloning modules between plant and bacterial systems, streamlining the workflow for recombinant protein production, especially in plant research.

## Introduction

Protein expression and purification are a basic need for biological research. Besides other available heterologous expression systems, recombinant protein expression in *E. coli* is the predominant form of protein production. However, achieving high yields of functional or correctly folded protein of interest (POI) remains challenging, as individual proteins can have very specific properties, and expression modalities need to be adjusted for every POI. Besides optimization of the expression conditions, like temperature or expression timing, identifying the ideal affinity tag for high yield and purity is crucial. Therefore, screening POI fusions with different small affinity-purification tags (e.g., 6His, StrepII) or larger solubility-enhancing tags (e.g., maltose-binding protein (MBP), glutathione S-transferase (GST)) is often necessary (Hochuli *et al*., 1988; Schmidt and Skerra, 2007; di Guana *et al*., 1988; Smith and Johnson, 1988). However, single affinity tags are often not sufficient to reach the desired properties, and thus, tandem affinity purification (TAP) tag, with or without inclusion of specific protease cleavage sites for tobacco etch virus [TEV] or human rhinovirus 3C (HRV3C, PreScission-like) protease, can be useful (Rigaut *et al*., 1999; Parks *et al*., 1994; Walker *et al*., 1994; Schwartz *et al*., 2025).

Nevertheless, classical cloning techniques are a severe bottleneck for this multi-construct screening process, as they have low efficiency and throughput, especially for multiple fragments in various combinations. GoldenGate assembly and broader ModularCloning (MoClo) systems circumvent this and enable efficient modular combination of multiple fragments (Weber *et al*., 2011). GoldenGate cloning relies on Type IIS restriction enzymes (e.g., BsaI or BpiI), which cleave outside their asymmetric recognition sequence, allowing directional assembly of multiple DNA fragments with a single enzyme and one-step reaction (Engler *et al*., 2008).

In plant research, multiple such GoldenGate-based cloning systems have been introduced, for example, for CRISPR-/Cas9 applications, native protein purifications from plants, or for modular assembly of plant expression cassettes with different complexities (Weber *et al*., 2011; Gantner *et al*., 2018; Grützner *et al*., 2021; Schwartz *et al*., 2025; Lampropoulos *et al*., 2013). GreenGate cloning is a widely used GoldenGate variant, as it is optimized for plant research and characterized by its simplicity, relying only on BsaI/Eco31I (Lampropoulos *et al*., 2013). This makes avoiding unwanted restriction enzyme sites easier and allows efficient cloning with fewer DNA modifications. However, despite its wide use for cloning of plant expression vectors, this system was not extended or adapted for compatibility with other expression systems, like heterologous protein expression in *E. coli*. This would allow an even more efficient cloning and screening process, as individual modules can be used for different purposes and do not require the recloning of the same fragments into the other system.

Here, we present a novel, comprehensive vector set for recombinant protein production in *E. coli* that is compatible with the GreenGate cloning system, as it uses the same overhang syntax. This toolkit enables the rapid, single-step combination of three modules for N- or C-terminal fusion of POIs. It includes diverse affinity tags of different sizes or solubilization properties, tag combinations, and versions of these tags with or without a specific protease cleavage site. We demonstrate the usability and flexibility of this vector set, enabling fast and efficient cloning for recombinant protein production of plant proteins.

## Results and Discussion

Ensuring reliable expression of heterologous proteins in *E. coli*, we generated an expression vector based on the established pDEST-HisMBP vector, harbouring a strong isopropyl-β-D-thiogalactopyranoside (IPTG) inducible TAC-Promoter (Ptac) to control POI expression (Nallamsetty *et al*., 2005). We introduced BsaI/Eco31I cloning sites compatible with the GreenGate-Overhangs B and D, to allow combinations of three modules, N-Tag, CDS, and C-Tag. (Fig 1). For efficient cloning and elimination of negative clones, we included the GreenGate cassette, containing a ccdB and chloramphenicol resistance gene (Lampropoulos *et al*., 2013). For the selection of positive clones and the distinction from the entry vectors that harbour an ampicillin-resistance (AmpR), we replaced the pDEST AmpR cassette with a kanamycin resistance cassette.

**Figure 1:**
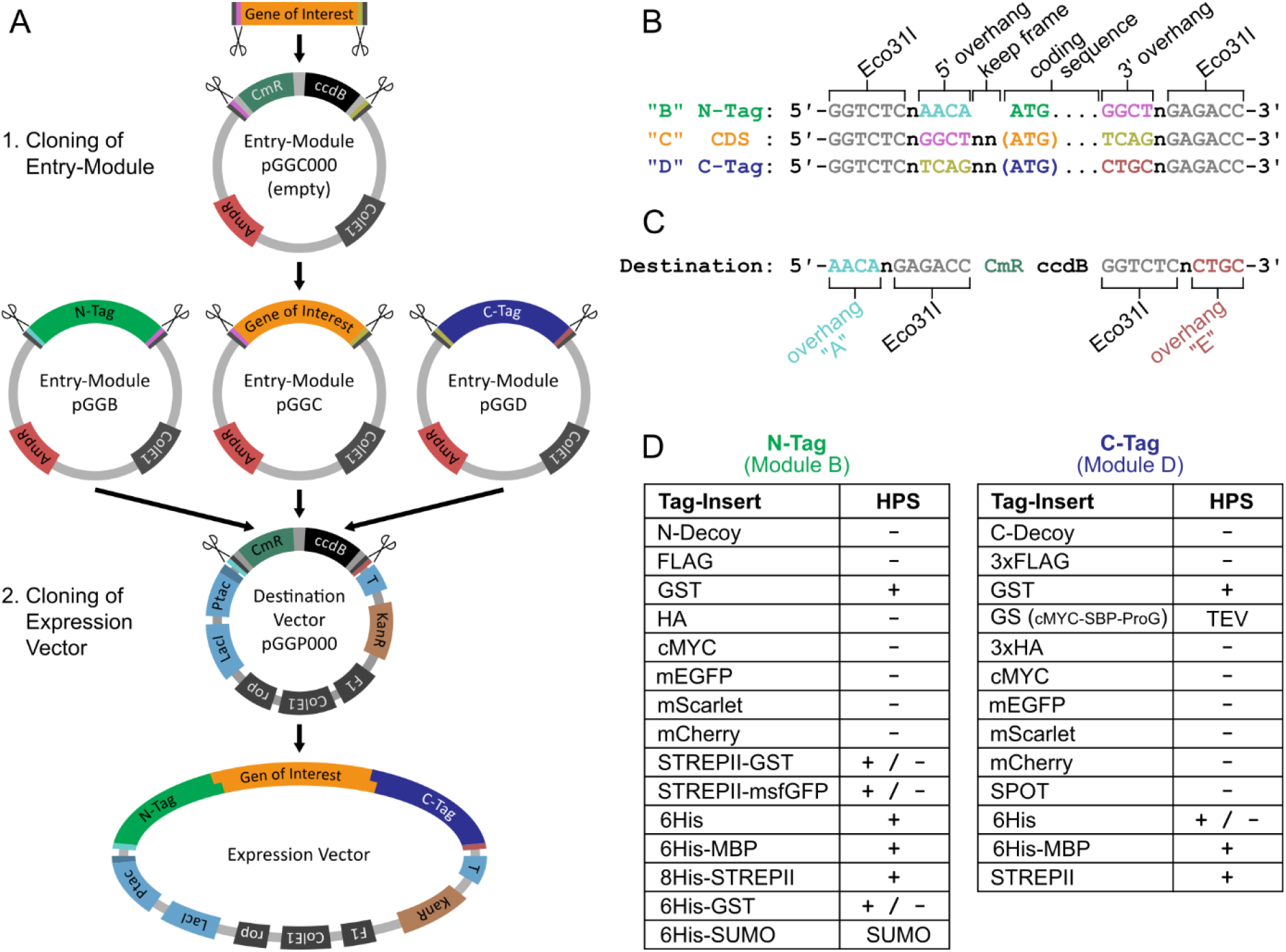
Cloning overview and features of the vector set. (**A**) Two-step cloning strategy of a gene of interest (GOI) using only BsaI/Eco31I-sites into an *E. coli* expression vector. Tags or GOI are first cloned into Enty-Modules (Ampicillin resistant (AmpR)) and then combined into a Destination-Vector (pGGP000, Kanamycin resistant (KanR)). (**B, C**) The used overhangs are compatible with the GreenGate-System and allow easy combination of three modules. Empty plasmids contain a chloramphenicol resistance for the specific selection during empty plasmid propagation and a ccdB gene for elimination of empty plasmids during cloning. (**D**) Precloned tag modules for N-terminal (Module B) or C-terminal (Module D) fusion, which are provided in the vector set. Presence of an HRV3C cleavage site (HPS) between tag and GOI is indicated.

Additionally, we generated a variety of tags in modules B and D for flexible addition of tags at both termini of the POI (Fig 1, Supplemental File 1). These tags include affinity-purification fragments, epitopes for immune detection, or combinations of both. These modules are available with or without HRV3C cleavage site (HPS) for cleavage of the tag from the POI (Walker *et al*., 1994). Assembly of an expression construct is possible in a single reaction, combining the destination-vector pGGP000 with tag modules and the GOI in module C or directly as a PCR product (Fig 1, and Material and Methods section for cloning instructions).

We generated multiple constructs to assess the expression and inducibility of various tags. These test constructs all contain a mEGFP combined with different combinations of tags. After transformation into Rosetta^™^(DE3) cells, cultures were induced with IPTG, and samples before and after induction were analysed by Western blot (Fig 2). All tested constructs showed little signal before induction and a strong signal after induction. However, in constructs including a GST-tag at the C-Terminus, the signal using anti-GFP antibodies was significantly reduced, even though full-length protein was sufficiently produced, shown by anti-GST antibody detection (Fig 2). In summary, this showed that independent of the tag, the fusion proteins are hardly produced before induction and strongly inducible with IPTG.

**Figure 2:**
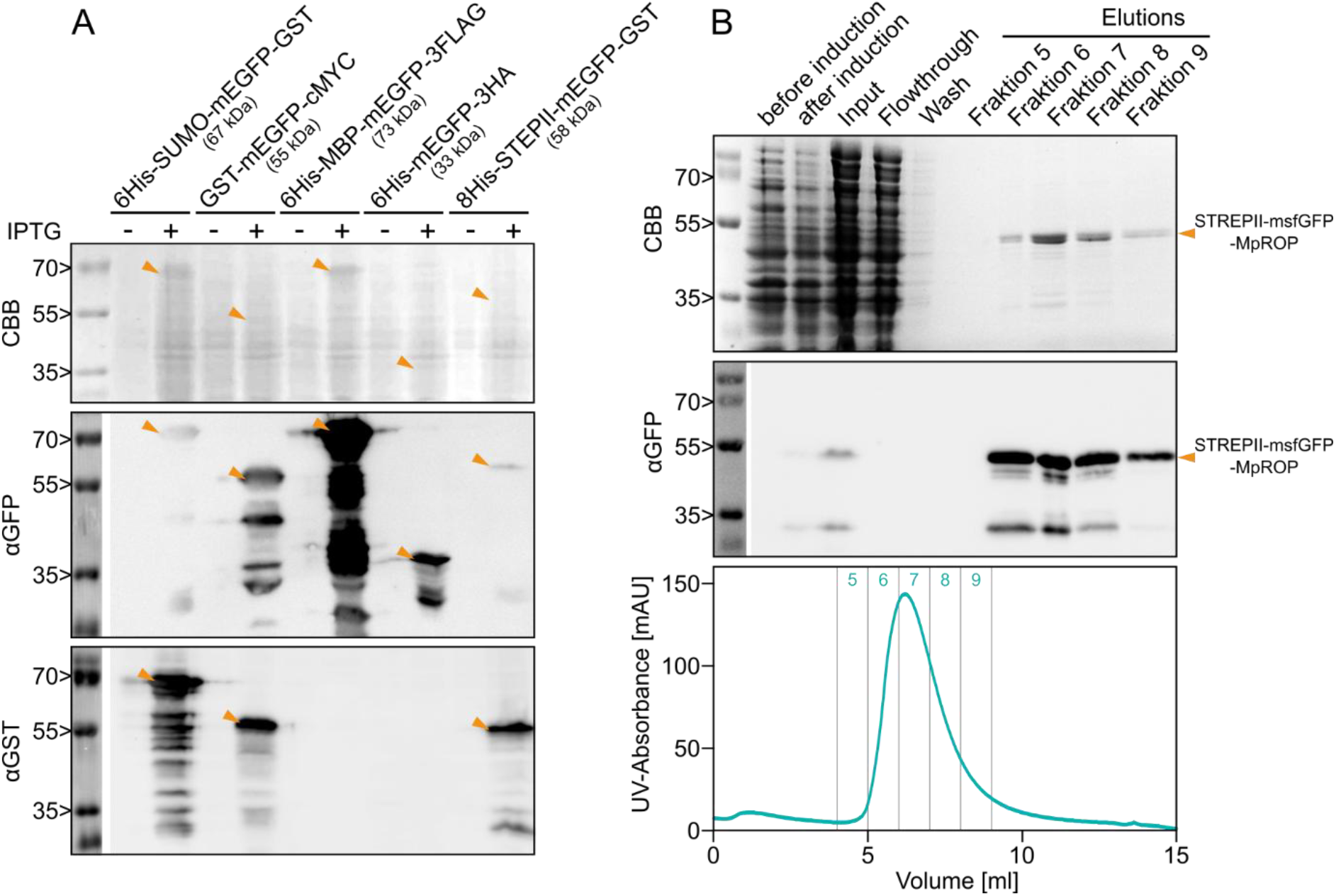
Example protein induction and purification. (**A**) Protein production of a variety of expression vectors that include multiple tags to test their usability. Samples before and 2 h after induction with IPTG are tested. Protein production is shown by Coomassie-stained SDS-PAGE (CBB), or by Western blots using antibodies against GFP (αGFP) or GST (αGST). Not all samples contain a GST tag and are expected to be empty. Sizes of the individual fusion proteins are indicated by arrowheads. (**B**) Example purification of a POI. STREPII-msfGFP-MpROP (50 kDa) was purified by FPLC and a Strep-Tactin®XT 4Flow® column. CBB-stained SDS-PAGE and αGFP Western blot show protein induction and purity after elution. The UV-trace of the FPLC elution is shown to indicate protein content in the elution fractions. Even though protein induction is low and not visible by CCB-staining, strong enrichment of the desired fusion protein is achieved.

To test the functionality of this system for affinity purification, we generated an expression construct of *Marchantia polymorpha* RHO OF PLANTS (MpROP), N-terminally tagged with a STREPII-msfGFP combination. After expression in Rosetta^™^(DE3) cells, we purified this fusion protein by Fast Protein Liquid Chromatography (FPLC) using a Strep-Tactin®XT 4Flow® column. Analysis by SDS-page and Coomassie blue staining or Western blot with anti-GFP antibody showed strong protein induction and high enrichment of POI (Fig 2). Even though the POI showed the strongest band at the expected 50 kDa, impurities were detectable, as is common in single-step affinity purifications.

To address this, we used the HRV3C protease, which is highly efficient in different buffer conditions and faster compared to the TEV protease, in combination with a tag that includes an HPS (Schwartz *et al*., 2025; Ullah *et al*., 2016). This allows the binding of the POI to the affinity matrix and a very specific release of the POI after washing (Fig 3A). We tested this strategy by tagging the *Marchantia polymorpha* PLANT-SPECIFIC ROP NUCLEOTIDE EXCHANGER domain (MpPRONE) of the ROP GUANINE NUCLEOTIDE EXCHANGE FACTOR (ROPGEF) protein with a 6His-GST TAP tag, separated by an HPS. After binding of the fusion protein to Glutathione-Sepharose 4B beads and removal of unspecific proteins, the sample was divided into two equal aliquots. MpPRONE was either specifically released by incubation with 6His-GST-HRV3C, or the 6His-GST-HPS-MpPRONE fusion protein was eluted by 40mM reduced glutathione (GSH) (Fig 3B).

**Figure 3:**
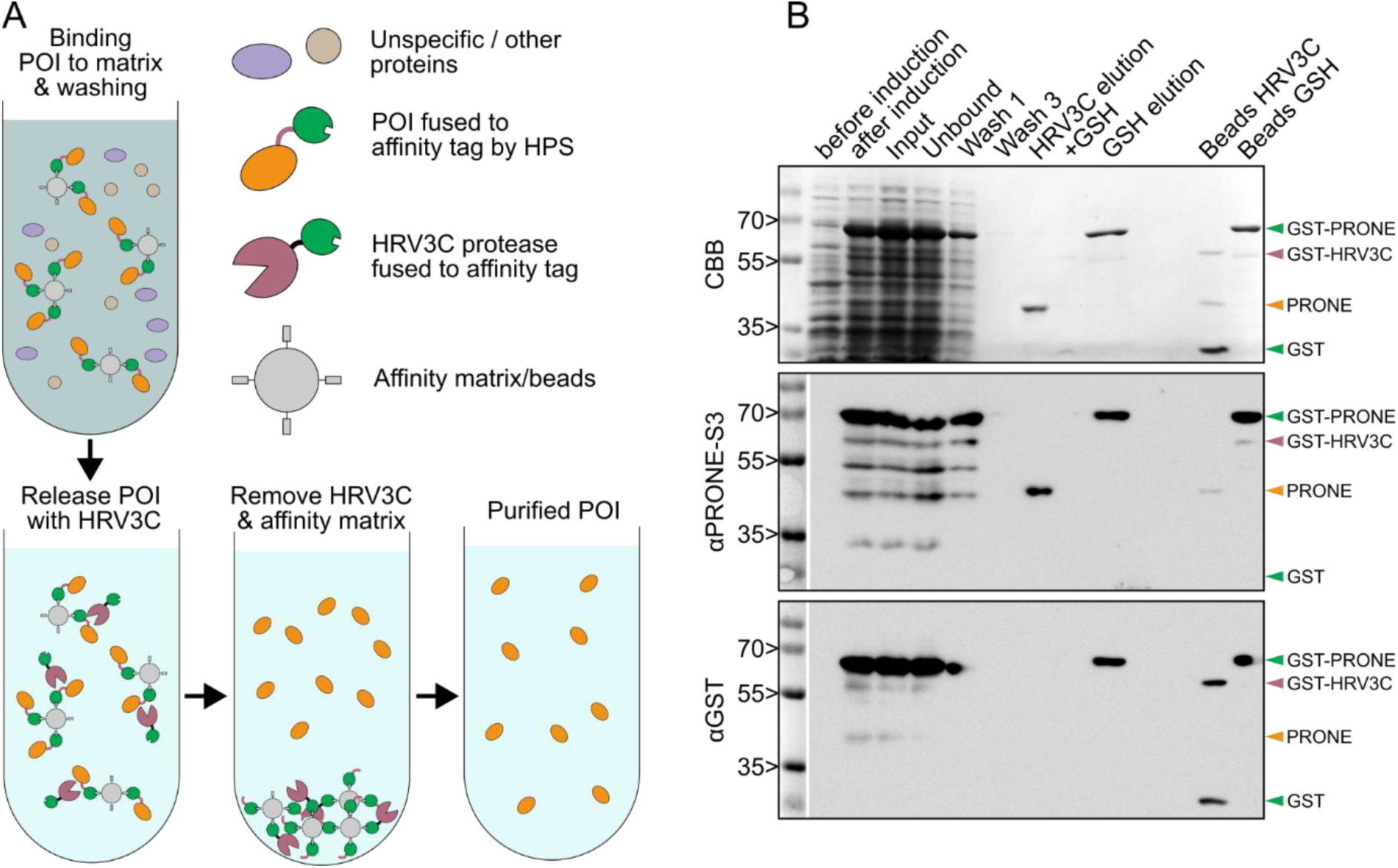
Tag-free protein purification using the HRV3C-Protease. (**A**) Schematic strategy of protein purification using an HRV3C-Protease (HRP). This strategy can be adapted with HRV3C-Protease fused to the corresponding affinity tag. First, the protein of interest (POI) fused to an affinity tag is bound to the affinity matrix. The POI is released from this complex by incubation with HRV3C-protease fused to the same affinity-tag as the POI. Thus, the protease also binds to the affinity matrix and can be separated from the POI. (**B**) Example purification of a POI. 6His-GST-HPS-MpPRONE. CBB-stained SDS-PAGE and αPRONE-S3 or αGST Western blots show protein induction and purity after elution. Before elution, the affinity-matrix beads were split and incubated either with HRV3C-Protease or 40mM reduced glutathione (GSH) to show the efficiency and resulting purity of both methods side by side. In the CBB-stained gel, additional bands can be observed in GSH eluted samples, while HRV3C released POI appear without these impurities. Additionally, samples of beads after elution show remaining fusion protein bound to the affinity matrix after elution with GSH. The size of the 6His-GST-HPS-MpPRONE fusion protein (70.4 kDa), released MpPRONE (42.7 kDa), free 6His-GST (27.7 kDa), and 6His-GST-HRV3C (61.6 kDa) is indicated.

Release of the POI by HRV3C resulted in higher purity compared to the GSH-eluted sample. Additionally, release of the POI by HRV3C had a higher efficiency in eluting the protein from the affinity matrix, as no POI remained bound to the beads compared to GSH-elution. Moreover, most of the protease is bound to the used beads, even though slight contaminations can reside. In this case, an incubation with additional Glutathione-Sepharose 4B beads can remove excess protease or contamination with full-length protein. This strategy provided a high purity of the POI and allows follow-up experiments without the negative influence of the tag.

In summary, this vector set allows efficient cloning of multiple tag combinations to a POI, especially in combination with existing GreenGate modules, which allows fast testing of different tags for affinity purifications or pull-down experiments. Moreover, the possible removal of tags by self-made HRV3C protease decreases impurities at low costs, providing pure POI for further investigations.

## Material and Methods

### Molecular cloning

To generate a GreenGate-compatible IPTG-inducible destination vector conferring kanamycin resistance, we used pDEST-HisMBP (AddGene #11085) as the backbone template (Nallamsetty *et al*., 2005). The ampicillin resistance cassette was exchanged for the kanamycin resistance cassette of the vector pGGX000, and a BsaI/Eco31I site in the rrnB terminator was mutated (Bouatta *et al*., 2025). Furthermore, the sequences downstream of the Tac promoter to the rrnB terminator were replaced with a CmR and ccdB-containing selection cassette with B and E overhangs from pGGX000. Plasmid selection and propagation were done in ccdB-resistant DB3.1 *E. coli*.

Entry modules were cloned according to the GreenGate system as described before (Bouatta *et al*., 2025; Lampropoulos *et al*., 2013). The fragments of interest were PCR-amplified with primers including the corresponding overhangs and BsaI sites, or synthesized (https://www.twistbioscience.com/de/), including the corresponding restriction sites. PCR product was purified from agarose gels, eluted with 45µl H_2_O. Synthesized DNA fragments or purified PCR products were digested together with 100 ng plasmid in the appropriate digest buffer for 30 min at 37 °C. After purification with a PCR-Purification Kit and elution in 20 µl, DNA was ligated with 1µl T4 DNA ligase (5 U/µl Thermo Scientific™) for 1h at room Temperature. For the entry modules including the HRV3C-Protease (pGPC005 and pGPC006), HRV3C fused to the NT* solubility tag was adapted from pET-NT-HRV3CP (Addgene #162795) (Abdelkader and Otting, 2021). For entry modules containing a monomeric superfolder GFP (msfGFP), we modified sfGFP (FPbase ID: B4SOW, Q80 variant) with a V206K mutation to enhance monomeric properties (Pédelacq *et al*., 2006; Zacharias *et al*., 2002).

Expression constructs were generated by combining 150 ng (100 ng/µl) of each entry module and 100ng pGGP000 (100ng/µl) with 1.5 µl Digest buffer, 1mM µl ATP, 1 µl HC T4 DNA ligase (30 U/µl Thermo Scientific™), and 1 µl Eco31I in a total volume of 15 µl. In a PCR cycler, 30 cycles of 37 °C for 2min and 16 °C for 3 min were performed, followed by digestion of undesired plasmids (50 °C, 10 min) and enzyme inactivation (85 °C, 10 min).

Entry and Destination reactions were transformed into ccdB-sensitive E. coli (e.g., DH5α) and plated on selection medium containing 100 µg/µL ampicillin (entry vectors) or 50 µg/µL kanamycin (destination vectors). For propagation of empty plasmids, ccdB-resistant DB3.1 *E. coli* were used.

All generated plasmids were sequenced to control all PCR-amplified fragments and ligation connections. All sequences of the used and generated plasmids are provided as .gb files in Supplemental File 2. For easy and cost-efficient distribution, the plasmids will be deposited in the European Plasmid Repository (https://www.plasmids.eu/) as a single Kit.

### Protein expression

A 4 mL overnight culture of LB medium containing 50 µg/µL kanamycin, 25 µg/mL chloramphenicol, and 1%°Glucose was inoculated with a colony of transformed Rosetta^™^(DE3) cells. 200 mL of the above medium was inoculated with 4 ml overnight culture and incubated to an OD_600_ of 0.6-0.8. Protein expression was induced with 1mM IPTG and incubated for 16h at 18 °C (or 3-4 h at 37° C) at 200 rpm. Cells were harvested (5 min, 4000 g, 4 °C), the cell pellet was washed with PBS, and stored at -20 °C.

### Protein purification

Cell pellets of 200ml culture (OD_600_ ∼3) were thawed on ice and resuspended in 20 ml lysis buffer (BasisBufferG: 50 mM Tris/HCl (pH 7.4), 150 mM NaCl, 1 mM EDTA (pH 8.3), 1 mM DTT, with freshly added 0.5% TritonX, 1mg/ml Lysozyme, 5 µg/ml DNase I and ½ Tablet of protease inhibitor cocktail (cOmplete™ Roche)). Cells were lysed by sonication on wet ice twice for 10 min, the lysate was cleared (10 min, 15.000 g, 4 °C), and the supernatant was used for purification.

100µl Glutathione Agarose 4B beads (Protino) were equilibrated in Lysis buffer, and GST-tagged proteins were bound to those beads by rotating for 2h at 4 °C. Unspecific proteins were removed by washing once with BasisBufferG containing 0.1% TritonX and twice with BasisBufferG containing 0.01% TritonX. POI was eluted with either 150µl 50mM Tris/HCl (pH 7.4) containing 40mM reduced glutathione (GSH), or by rotating incubation in 150 µl BasisBufferG containing 5-10 µl purified His-GST-HRV3C protease (pGGP001) for 16-20h at 4 °C and removal of the beads. In case of small contaminations with full-length protein or HRV3C protease, which needs to be removed, the supernatant was incubated with 50 µl Glutathione Agarose 4B beads and rotated for 30 min at 4 °C.

For His-GST-HRV3C protease (pGGP001) purification, the protein of two pellets was eluted with 250 µl 50 mM Tris/HCl (pH 7.4) containing 25 mM reduced glutathione, dialysed with 500 ml 50 mM sodium phosphate buffer (pH 7.2) containing 150 mM NaCl and 5% glycerol, and stored at -20 °C in 50 µl Aliquots after adding Glycerol to 20%.

StrepII-tagged proteins were filtered, degassed, and purified using FPLC (Cytiva ÄKTA pure™) in a 1 ml Strep-Tactin®XT 4Flow® column (IBA Lifesciences). Unspecific proteins were removed by washing with 15 ml BasisBufferS: 100 mM Tris/HCl (pH 8.0), 150 mM NaCl, 1 mM EDTA. POIs were eluted with 14 ml BasisBufferS containing 25 mM biotin. Columns were regenerated with 15 mL 3 M MgCl_2_.

### Protein detection

Protein expression and purification were controlled on Coomassie-stained SDS-PAGE gels, or by Western blot using antibodies in 5% milk in TBS-T for GST detection (1:5000 αGST (goat, Cytiva 27-4577-01) and 1:5000 αGoat-HRP (rabbit, Sigma-Aldrich A5420)) and for GFP detection (1:1000 αGFP (mouse, Roche 11814460001) and 1:1000 αMouse-HRP (rabbit, Sigma-Aldrich A9044)). For PRONE domain detection, we used a custom rabbit antiserum (1:1000) against the partially conserved peptide NFRGHNEFWYVSRDSEEG within the PRONE domain of AtROPGEF8. This Serum detected all tested PRONE domains from *Arabidopsis* and *Marchantia*. As secondary antibody we used 1:1000 αRabbit-HRP (goat, Sigma-Aldrich A0545)

## Supporting information

Supplemental File 1_VectorTable

Supplemental File 2_Vectors

## Acknowledgement

We thank Jan Lohmann for providing us with the GreenGate vector set ahead of publication, and Ingo Heilmann for a plasmid containing sfGFP. Additionally, we thank Claus Schwechheimer and Thomas Dresselhaus for providing access to their facilities and the FPLC instruments.

## Author contributions

AB and PD conceived the project. AB, AL, FA, MDLB, and PD generated material. MDLB, FA, and AL established the experimental setup and conducted experiments. PD compiled the figures and wrote the manuscript with MDLB and AB, with input from AL and FA.

## Supplemental Files

### Supplemental File 1: List of generated and used vectors

Vector list, including descriptions, ID, and position within the deposited Kit. All listed vectors are deposited in the European Plasmid Repository (https://www.plasmids.eu/).

### Supplemental file 2: Vector sequences

All vector sequences, including annotation of the functional domains, are provided as .gb files and compiled in a single .zip folder.

## References

Abdelkader, E.H. and Otting, G. (2021) NT*-HRV3CP: An optimized construct of human rhinovirus 14 3C protease for high-yield expression and fast affinity-tag cleavage. Journal of Biotechnology, 325, 145–151.

Bouatta, A.M., Anzenberger, F., Riederauer, L., Lepper, A. and Denninger, P. (2025) Polarized subcellular activation of Rho proteins by specific ROPGEFs drives pollen germination in Arabidopsis thaliana. PLOS Biology, 23, e3003139.

Engler, C., Kandzia, R. and Marillonnet, S. (2008) A One Pot, One Step, Precision Cloning Method with High Throughput Capability. PLOS ONE, 3, e3647.

Gantner, J., Ordon, J., Ilse, T., et al. (2018) Peripheral infrastructure vectors and an extended set of plant parts for the Modular Cloning system. PLOS ONE, 13, e0197185.

Grützner, R., Martin, P., Horn, C., Mortensen, S., Cram, E.J., Lee-Parsons, C.W.T., Stuttmann, J. and Marillonnet, S. (2021) High-efficiency genome editing in plants mediated by a Cas9 gene containing multiple introns. Plant Communications, 2, 100135.

Guana, C. di, Lib, P., Riggsa, P.D. and Inouyeb, H. (1988) Vectors that facilitate the expression and purification of foreign peptides in Escherichia coli by fusion to maltose-binding protein. Gene, 67, 21–30.

Hochuli, E., Bannwarth, W., Döbeli, H., Gentz, R. and Stüber, D. (1988) Genetic Approach to Facilitate Purification of Recombinant Proteins with a Novel Metal Chelate Adsorbent. Nat Biotechnol, 6, 1321–1325.

Lampropoulos, A., Sutikovic, Z., Wenzl, C., Maegele, I., Lohmann, J.U. and Forner, J. (2013) GreenGate -A novel, versatile, and efficient cloning system for plant transgenesis P. J. Janssen, ed. PLoS ONE, 8, e83043.

Nallamsetty, S., Austin, B.P., Penrose, K.J. and Waugh, D.S. (2005) Gateway vectors for the production of combinatorially-tagged His6-MBP fusion proteins in the cytoplasm and periplasm of Escherichia coli. Protein Science, 14, 2964–2971.

Parks, T.D., Leuther, K.K., Howard, E.D., Johnston, S.A. and Dougherty, W.G. (1994) Release of Proteins and Peptides from Fusion Proteins Using a Recombinant Plant Virus Proteinase. Analytical Biochemistry, 216, 413–417.

Pédelacq, J.-D., Cabantous, S., Tran, T., Terwilliger, T.C. and Waldo, G.S. (2006) Engineering and characterization of a superfolder green fluorescent protein. Nat Biotechnol, 24, 79–88.

Rigaut, G., Shevchenko, A., Rutz, B., Wilm, M., Mann, M. and Séraphin, B. (1999) A generic protein purification method for protein complex characterization and proteome exploration. Nat Biotechnol, 17, 1030–1032.

Schmidt, T.G. and Skerra, A. (2007) The Strep-tag system for one-step purification and high-affinity detection or capturing of proteins. Nat Protoc, 2, 1528–1535.

Schwartz, S., Engstler, C., Mühlbauer, S., Windenbach, E., Wunder, T., Kunz, H.-H. and Brandt, B. (2025) Catch & Release—rapid cost-effective protein purification from plants using a DIY GFP-Trap-protease approach. The Plant Journal, 124, e70544.

Smith, D.B. and Johnson, K.S. (1988) Single-step purification of polypeptides expressed in Escherichia coli as fusions with glutathione S-transferase. Gene, 67, 31–40.

Ullah, R., Shah, M.A., Tufail, S., Ismat, F., Imran, M., Iqbal, M., Mirza, O. and Rhaman, M. (2016) Activity of the Human Rhinovirus 3C Protease Studied in Various Buffers, Additives and Detergents Solutions for Recombinant Protein Production. PLOS ONE, 11, e0153436.

Walker, P.A., Leong, L.E.-C., Ng, P.W.P., Tan, S.H., Waller, S., Murphy, D. and Porter, A.G. (1994) Efficient and Rapid Affinity Purification of Proteins Using Recombinant Fusion Proteases. Nat Biotechnol, 12, 601–605.

Weber, E., Engler, C., Gruetzner, R., Werner, S. and Marillonnet, S. (2011) A Modular Cloning System for Standardized Assembly of Multigene Constructs. PLOS ONE, 6, e16765.

Zacharias, D.A., Violin, J.D., Newton, A.C. and Tsien, R.Y. (2002) Partitioning of Lipid-Modified Monomeric GFPs into Membrane Microdomains of Live Cells. Science, 296, 913–916.

